# Reproducible functional connectivity alterations are associated with autism spectrum disorder

**DOI:** 10.1101/303115

**Authors:** Štefan Holiga, Joerg F. Hipp, Christopher H. Chatham, Pilar Garces, Will Spooren, Xavier Liogier D’Ardhuy, Alessandro Bertolino, Céline Bouquet, Jan K Buitelaar, Carsten Bours, Annika Rausch, Marianne Oldehinkel, Manuel Bouvard, Anouck Amestoy, Mireille Caralp, Sonia Gueguen, Myriam Ly-Le Moal, Josselin Houenou, Christian F. Beckmann, Eva Loth, Declan Murphy, Tony Charman, Julian Tillmann, Charles Laidi, Richard Delorme, Anita Beggiato, Alexandru Gaman, Isabelle Scheid, Marion Leboyer, Marc-Antoine d’Albis, Jeff Sevigny, Christian Czech, Federico Bolognani, Garry D. Honey, Juergen Dukart

## Abstract

Despite the high clinical burden little is known about pathophysiology underlying autism spectrum disorder (ASD). Recent resting state functional magnetic resonance imaging (rs-fMRI) studies have found atypical synchronization of brain activity in ASD. However, no consensus has been reached on the nature and clinical relevance of these alterations. Here we address these questions in the most comprehensive, large-scale effort to date comprising evaluation of four large ASD cohorts. We followed a strict exploration and replication procedure to identify core rs-fMRI functional connectivity (degree centrality) alterations associated with ASD as compared to typically developing (TD) controls (ASD: *N*=841, TD: *N*=984). We then tested for associations of these imaging phenotypes with clinical and demographic factors such as age, sex, medication status and clinical symptom severity. We find reproducible patterns of ASD-associated functional hyper- and hypo-connectivity with hypo-connectivity being primarily restricted to sensory-motor regions and hyper-connectivity hubs being predominately located in prefrontal and parietal cortices. We establish shifts in between-network connectivity from outside to within the identified regions as a key driver of these abnormalities. The magnitude of these alterations is linked to core ASD symptoms related to communication and social interaction and is not affected by age, sex or medication status. The identified brain functional alterations provide a reproducible pathophysiological phenotype underlying the diagnosis of ASD reconciling previous divergent findings. The large effect sizes in standardized cohorts and the link to clinical symptoms emphasize the importance of the identified imaging alterations as potential treatment and stratification biomarkers for ASD.

## Introduction

Autism spectrum disorder (ASD) is a neurodevelopmental disorder characterized by deficits in social interaction and communication, and by the presence of restricted and stereotyped behaviors and sensory anomalies*(1)*. ASD occurs in up to about 1% of general child population*(2)* and despite its early-childhood onset and lifelong persistence, surprisingly little is known about the neurobiology underlying its cause and pathogenesis*(3)*. Based on various divergent findings several theories have been proposed to explain ASD at genetic, neuropathology, systemic and behavioral levels*(4, 5)*. On a systemic level, the hypothesis of abnormal brain functional connectivity has been proposed more than a decade ago*(6)*.

One of the prevailing experimental approaches to study brain functional connectivity non-invasively is by resting-state functional magnetic resonance imaging (rs-fMRI). Rs-fMRI reflects spontaneous neuronal activity by detecting slow fluctuations of local oxygen demands. Inter-regional temporal synchronization of the intrinsic hemodynamics is then considered as an index of functional connectivity*(7)*. A significant body of evidence now supports the existence of such functional connectivity alterations in ASD*(8–14)*. However, the field has still failed to converge on the anatomy, nature, directionality and clinical relevance of the observed alterations. While some reports points to an under-connectivity phenotype*(15–17)*, others reported the opposite*(11, 18)*, and some observed a mix of both phenomena*(9, 19)*. Besides inherent disease heterogeneity, this discordance has been proposed to arise from age or methodological differences between studies*(17, 20, 21)*, or failures to account for medication effects in the analyses*(22)*. Yet other work highlights the importance of spatial variability in connectivity as a key source of heterogeneity in ASD*(19, 23)*. A common caveat in interpretation of these studies is the lack of replication of respective findings. A consensus on the nature and clinical relevance of the functional connectivity changes in ASD therefore still needs to be reached. These uncertainties limit the usability of functional connectivity as a potential diagnostic or treatment biomarker for ASD.

Here we aimed to address these inconsistent findings by performing a large-scale evaluation and characterization of functional connectivity alterations in ASD in the most comprehensive effort to date comprising a rigorous replication strategy in four independent cohorts. More specifically, we used the EU-AIMS LEAP dataset for exploration and the other three cohorts (ABIDE I, ABIDE II and InFoR) for replication. We further evaluated the link between these alterations and age, medication status and clinical symptoms.

## Results

### Clinical and demographic characteristics

Exclusion of individuals with low IQ and removal of poor image quality data resulted in exclusion of 35, 39, 33 and 14% of available EU-AIMS LEAP, ABIDE I, ABIDE II and InFoR data, respectively (s. Table 1 for final N). For all cohorts, mean age did not differ between ASD patients and TD and was comparable between EU-AIMS LEAP and ABIDE I (Table 1). ABIDE II cohort was significantly younger (all p<.001) and the InFoR only comprised adult ASD patients. A significantly higher fraction of males was observed in the ABIDE I and II as compared to EU-AIMS LEAP and InFoR (for ABIDE I). ASD patients in all cohorts had average level IQ although significantly lower than that for TD in all cohorts except InFoR (Table 1).

**Table 1.**
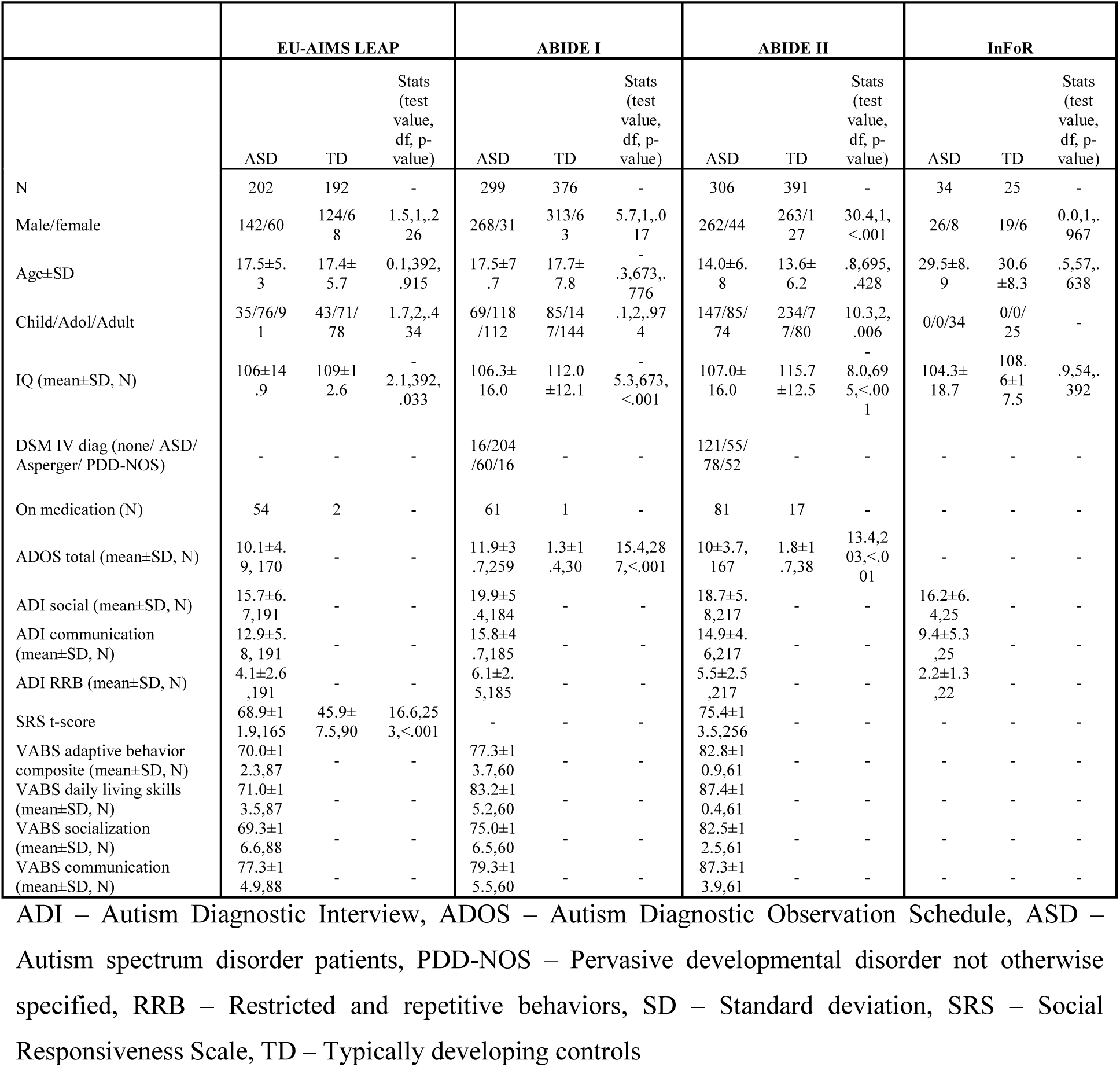
Clinical and demographic characteristics

### Reproducible degree centrality alterations in ASD patients

To test for functional connectivity differences between ASD patients and TD, we used the EU-AIMS LEAP dataset for exploratory analyses and the other three cohorts for replication. Degree centrality (DC) was chosen as an unbiased count-based functional connectivity measure that assigns to each voxel the sum of all correlation coefficients between the time series of that voxel and all other voxels in the brain exceeding a prespecified threshold. Group comparisons in the EU-AIMS cohort revealed significant DC increases in ASD patients in two clusters comprising bilateral parietal, prefrontal and anterior and posterior cingulate regions (Figure 1a and Table S8). Significantly decreased DC was observed in a cluster covering bilateral primary sensory motor cortices and right temporal regions, insula, amygdala and hippocampus. These results remained significant and qualitatively and quantitatively highly similar to the initial findings after controlling for between-subject differences in motion (Figure S2).

**Figure 1.**
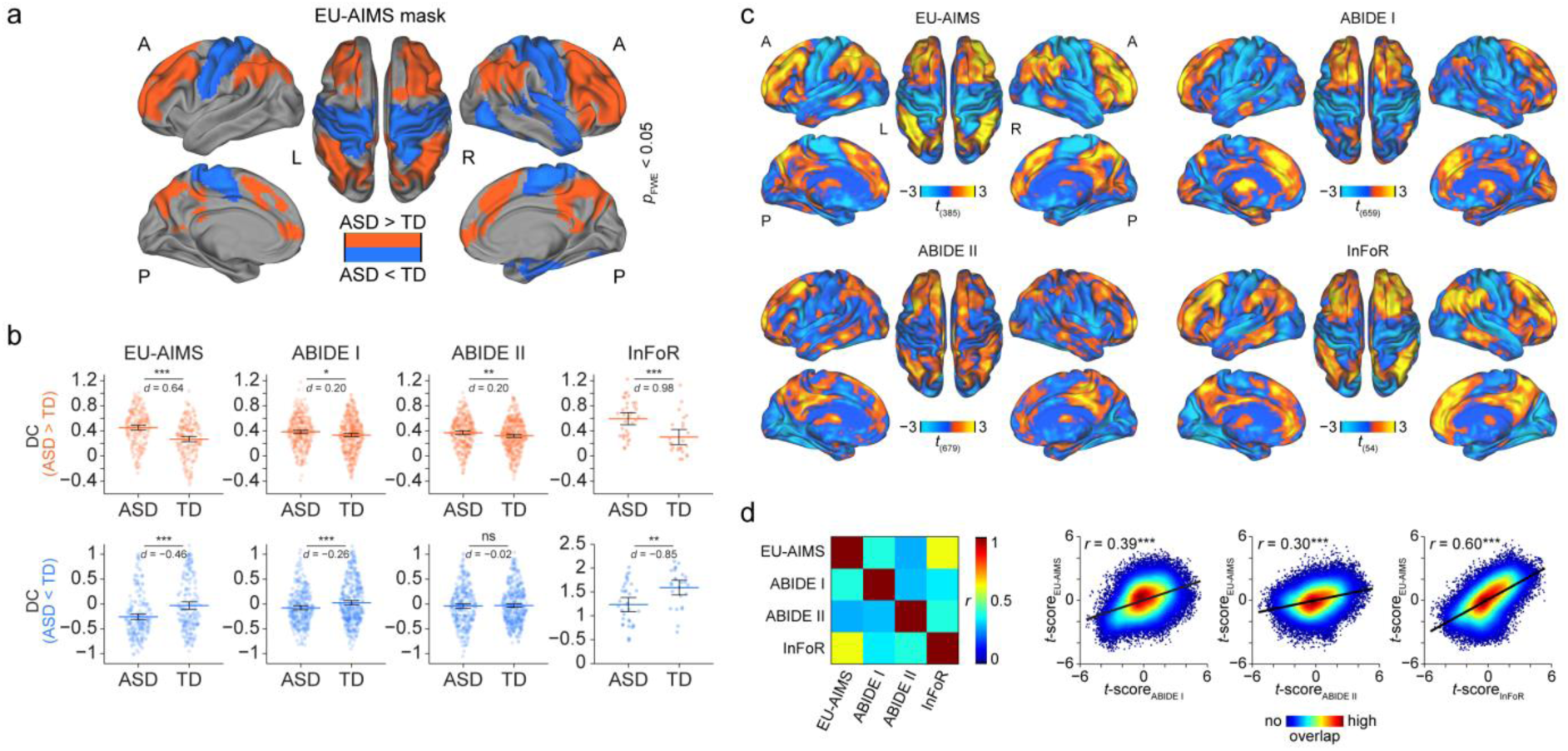
Outcomes of functional connectivity comparisons between ASD patients and TD controls. **(a)** Regions showing significant DC increases (red) and decreases (blue) in ASD patients in the reference EU-AIMS LEAP cohort. **(b)** Distribution of DC values (first eigenvariate over the masks’ voxels) across subjects for the hyper-connected (top row, red) and hypo-connected (bottom row, blue) network separately for ASD patients and TD controls. **(c)** Unthresholded *t*-maps showing regions with increased (warm colors) and decreased (cold colors) DC in ASD patients as compared to TD controls. **(d)** Spatial correspondence of the *t*-maps between the reference data set (EU-AIMS LEAP) and all remaining replication data sets (ABIDE I, ABIDE II, InFoR) in form of a pair-wise correlation matrix and individual correlation plots. A – anterior, ASD – autism spectrum disorder, d – Cohen’s d effect size, DC – degree centrality, FWE – family-wise error, L – left, P – posterior, R – right, TD – typically developing. \**p* < 0.05, \*\**p* < 0.01, \*\*\**p* < 0.001.

DC extracted from ABIDE I, ABIDE II and InFoR based on the EU-AIMS LEAP DC increase mask revealed significant increases in ASD patients in all three replication cohorts (Figure 1b). Conversely, the DC decreases were successfully replicated in two out of three cohorts (ABIDE I and InFoR). Evaluation of spatial similarity of the unthresholded DC alteration patterns revealed highly significant correlations of the ASD profiles across all four cohorts (all *p*<0.001, Figure 1c and d).

### Degree centrality alterations in ASD are driven by shifts in connectivity

Having identified replicable DC alterations in ASD patients we aimed to explore the nature of these connectivity differences. To this end, we compared the whole-brain histograms of correlation coefficients observed in ASD patients to TD revealing a nearly identical distribution of correlation strength (Figure S1). We next computed effect sizes for differentiation between ASD patients and TD for four types of metrics comparing means, variances, proportions and shifts in connectivity within, outside and from within to outside the identified DC alteration clusters (Figure 2a, b). These analyses revealed that shifts in correlations from outside to inside the DC increase mask showed the largest effect size (Figure 2b). This large effect reflects that in ASD voxels falling outside the DC increase mask show reduced connectivity to one another and an increased connectivity to voxels within the mask. This hyper-connectivity index was closely negatively correlated with the initial DC finding (*r*=−.72; *p*<.001; N=394).

**Figure 2.**
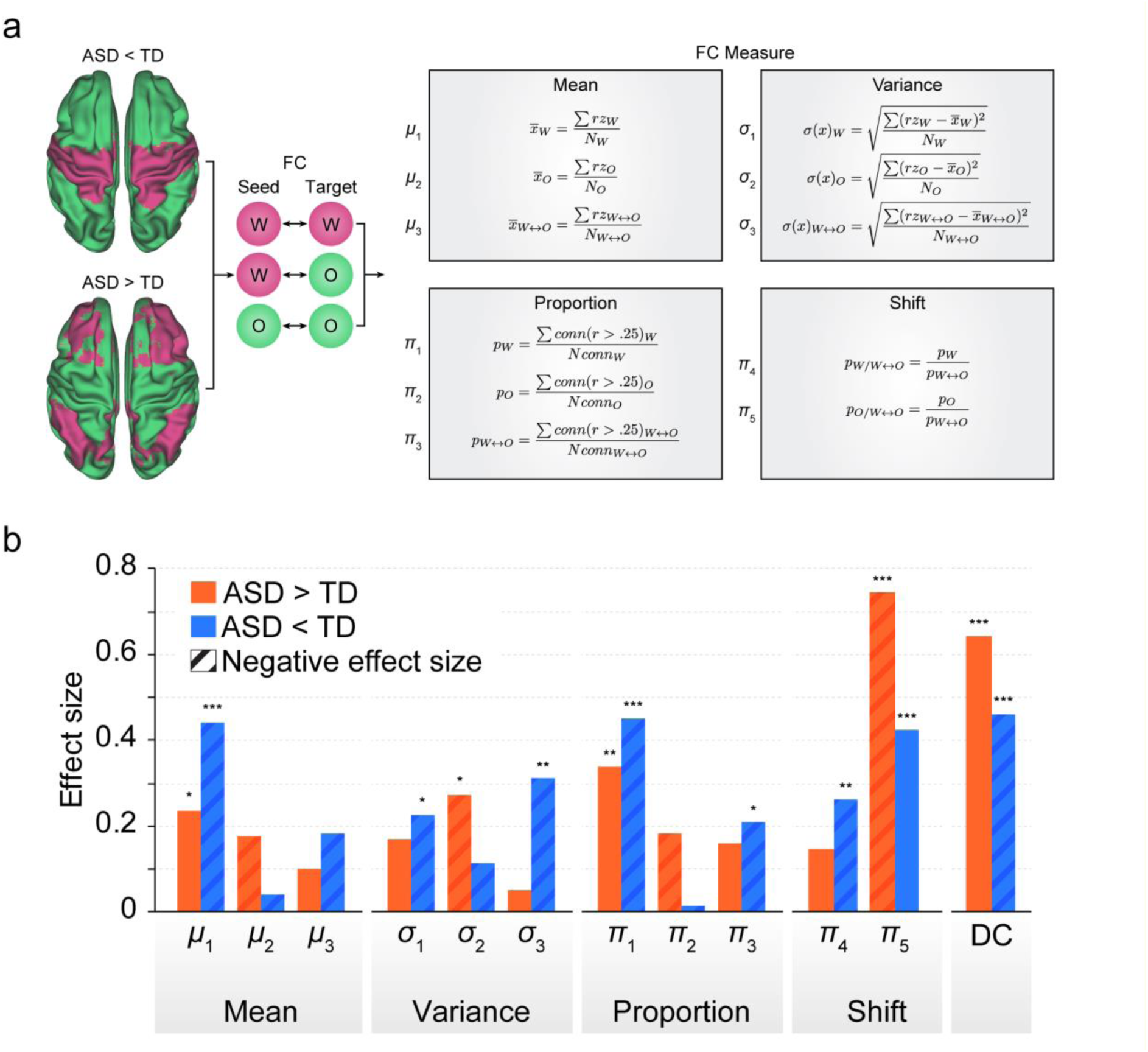
Functional connectivity indices and their effect size. **(a)** Schematic representation of different combinations of seed regions and search spaces used for functional connectivity analyses along with formulas used for calculation of the respective functional connectivity indices. **(b)** Effect sizes (absolute value – for comparability purpose) of the functional connectivity indices based on degree centrality masks for the contrasts ASD > TD and TD > ASD. Background colors reflect the four types of metrics described in (a). ASD – autism spectrum disorder, DC – degree centrality, o – outside, *r* – correlation coefficient, *rz* – Fisher’s z-transformed correlation coefficient, TD – Typically developing, w – within, prop – proportion, Std – standard deviation. \**p* < 0.05, \*\**p* < 0.01, \*\*\**p* < 0.001, - – indicates the original negative sign of the respective index.

For the DC decrease findings, three indices revealed similar effect size as the initial DC finding: reduced mean connectivity and number of connected voxels within the DC decrease mask, and shift in connectivity from outside to inside the DC decrease mask. For the latter, the proportion of voxels connected to each other outside the DC decrease mask increased and the proportion of voxels connected from outside to inside decreased. Very strong correlations were observed between all these indices and the initial DC decrease findings (mean connectivity: *r*=.81; *p*<.001, proportion of connected voxels: *r*=.84; *p*<.001, shift in connectivity: *r*=−.85; *p*<.001; N=394). As the shift in connectivity showed the strongest correlation with the initial DC findings, this hypo-connectivity index was selected for further testing of associations with clinical symptoms.

The further exploration in the EU-AIMS LEAP dataset of the choice of correlational threshold for DC computation on the observed differentiation between ASD and TD revealed a maximum differentiation for the hyper- and hypo-connectivity index at r=0.15 and r=0.4, respectively (Figure 3c). Importantly, for both indices the effect size for differentiation between ASD and TD obtained at the initially chosen DC threshold (r=0.25) was very close to the maximum observed with those optimized thresholds.

**Figure 3.**
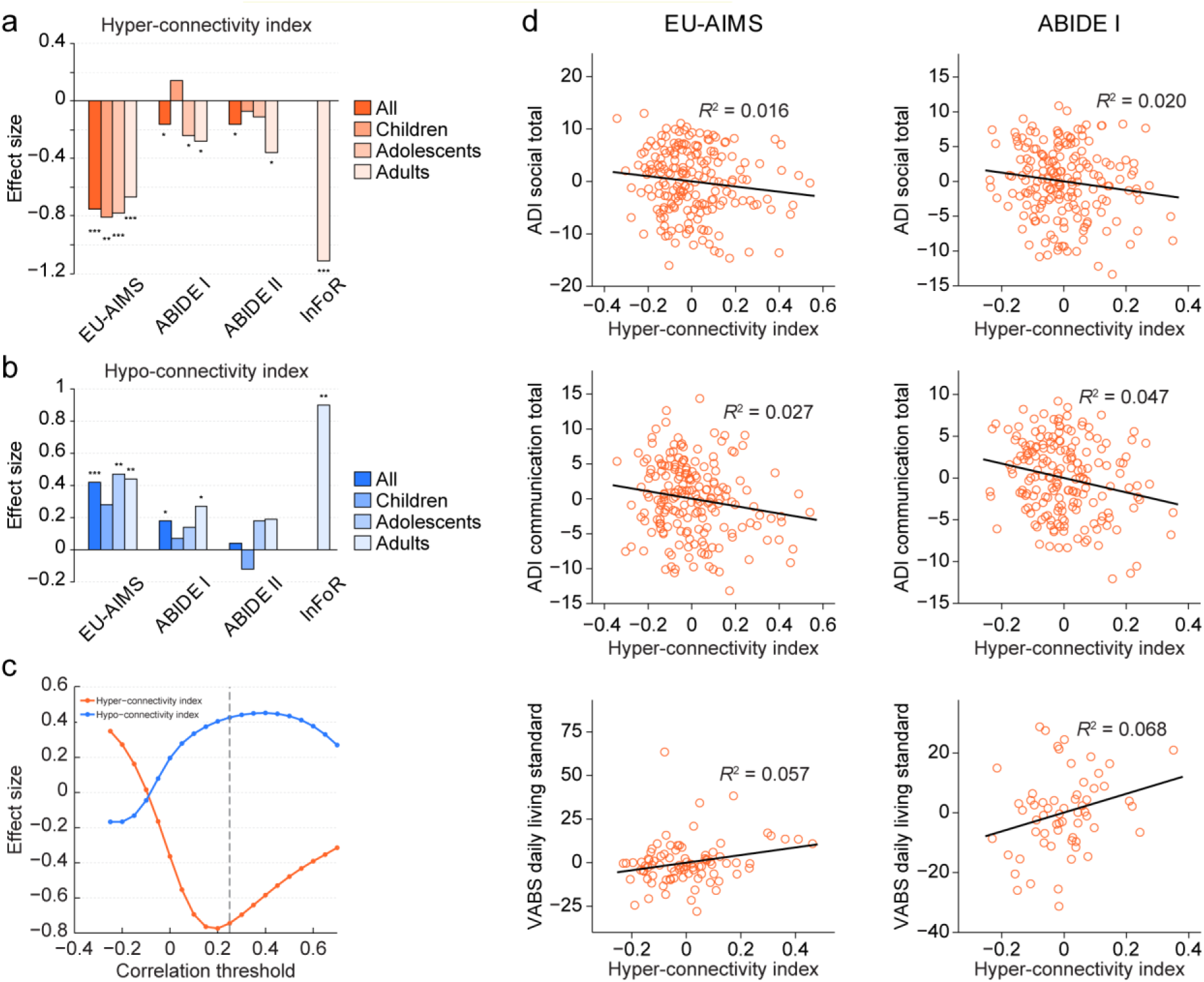
Relationships between hyper- and hypo-connectivity indices, age and clinical outcomes. **(a)** Effect size of the *hyper-connectivity index* for all patients and split by age groups (children [<12 years], adolescents [12–18 years], adults [>18 years]) for contrast ASD > TD (red). **(b)** The equivalent of (a) for the hypo-connectivity index (blue). **(c)** Effect size for the *hyper- and hypo-connectivity indices* obtained in the EU-AIMS LEAP dataset plotted versus the applied correlational threshold. Dashed line indicates the correlational threshold used for the initial degree centrality computation **(d)** Significant correlations of the hyper-connectivity indices with ASD clinical scales in the EU-AIMS LEAP and ABIDE I data sets (adjusted for the effects of age, sex, site and IQ). ASD – autism spectrum disorder, comm – communication, dls – daily living skills, \**p* < 0.05, \*\**p* < 0.01, \*\*\**p* < 0.001.

### Connectivity differences are linked to demographic and clinical factors

We next evaluated if and how these hyper- and hypo-connectivity indices relate to clinical characteristics, age, sex and medication status in ASD patients. First, we tested if the hyper- and hypo-connectivity alterations persist when splitting the cohorts into children, adolescents and adults. For the hyper-connectivity index, significant differences and similar effect sizes were observed for all age groups in the EU-AIMS LEAP cohort (Figure 3a). In other cohorts significant differences between ASD patients and TD were observed for adolescents (ABIDE I only) and adults (ABIDE I and II) but not for children. The hypo-connectivity index showed consistently larger effect sizes in the adolescent and adult populations as compared to children in all three cohorts (Figure 3b). In formal testing for age category by diagnosis interactions on the hyper- and hypo-connectivity indices using analysis of variance (ANOVA) a marginally significant interaction was only observed for the hyper-connectivity index in ABIDE I (F(2,656)=2.4;p=.095) and the hypo-connectivity index in ABIDE 2 (F(2,676)=2.4;p=.095).

In analysis of variance (ANOVA) testing for the effects of sex on the observed hyper- and hypo-connectivity indices no significant sex-by-diagnosis group interactions were observed for any of the cohorts (all p>.1). A significant effect of psychotropic medication on functional connectivity was only found for DC increases in the EU-AIMS LEAP cohort with ASD patients on medication being closer to the TD population (Table S9).

Following a similar exploration and replication strategy as for group comparisons, GLMs were computed in the EU-AIMS LEAP dataset to evaluate if the obtained hyper- and hypo-connectivity indices are linked to clinical severity. These analyses revealed overall three significant or marginally significant relationships between the hyper-connectivity index and Autism Diagnostic Interview (ADI) social and communication subscales and Vineland Adaptive Behavior Scale (VABS, *(24)*) daily living skills standard score (Table 2 and Table S10). Stronger connectivity alterations were associated with stronger clinical impairment (Figure 3d). In replication analyses, all three relationships found in EU-AIMS LEAP were also significant or marginally significant in the ABIDE I cohort but not in the substantially younger ABIDE II subset (Table 2, Table S11).

**Table 2.** Results of correlations between hyper- and hypo-connectivity indices and clinical scales

| Scale | Connectivity measure | EU-AIMS LEAP | ABIDE I | ABIDE II |
| --- | --- | --- | --- | --- |
| ADI social total | hyper-conn | ***$F(1,182)=3; p=0.087$*** | ***$F(1,170)=3.5; p=0.065$*** | *$F(1,204)=0; p=0.963$* |
| ADI communication total | hyper-conn | ***$F(1,182)=5; p=0.026$*** | ***$F(1,171)=8.7; p=0.004$*** | *$F(1,204)=0.3; p=0.602$* |
| VABS daily living standard | hyper-conn | ***$F(1,80)=5.3; p=0.024$*** | ***$F(1,55)=4.1; p=0.048$*** | *$F(1,55)=0.1; p=0.75$* |

ADI – Autism Diagnostic Interview, ASD – Autism spectrum disorder, DC – Degree centrality, conn – connectivity index, VABS – Vineland Adaptive Behavior Scales, Significant or marginally significant relationships between functional connectivity indices and respective clinical scales are indicated in bold.

## Discussion

Here we provide for the first time evidence of reproducible group-level functional hyper- and hypo-connectivity alterations in ASD patients as demonstrated across several large single- and multi-center cohorts. We further show that these alterations persist when controlling for motion and medication effects and are linked to clinical symptoms as captured by several clinical and behavioral scales.

We reconcile previous divergent literature findings and report evidence of both functional hypo- and hyper-connectivity in ASD patients, with spatially-varying signatures that are consistent with recent single cohort evidence *(9, 11, 17, 18)*. The identified hypo- and hyper-connectivity patterns show significant spatial similarity across all cohorts with the hypo-connectivity being primarily restricted to sensory-motor regions and hyper-connectivity hubs being predominately located in prefrontal and parietal cortices. Both anatomical networks are consistent with some previous work reporting connectivity alterations in respective networks*(9, 19)*. Consistent with some previous reports, we find the overall global connectivity to be preserved in ASD*(19)*.

Anatomically, the DC increase regions closely correspond to the well-established central executive network*(25–27)*. This network has been closely implicated in high-level cognitive functions such as planning, decision making, and the control of attention and working memory*(26)*. All of these functions have been consistently shown to be affected in ASD*(28–30)*. Moreover, we demonstrate that the DC increases originate specifically from increased engagement of this network by regions outside the network. Similarly, DC decreases are mainly driven by disengagement of the sensory-motor network from other regions. However, we also observe a similar magnitude of reduced connectivity within the sensory-motor network suggesting that both processes contribute to DC decreases. Biologically, these connectivity shifts may support the idea that ASD patients are unable to engage or disengage specific networks to the extent of TD, and may therefore underlie deficits in mental switching and cognitive flexibility*(28, 29)*.

Estimates of effect size range from small to very large across cohorts. Importantly, the ABIDE I and II datasets are retrospectively established cohorts with large variation in diagnostic and inclusion criteria, sequence quality, scanning duration and other parameters such as resting state acquisition instructions. This variability may explain the observed low effect sizes. In contrast, the standardized multi-center EU-AIMS and the single center InFoR cohort show a large effect size for the hyper-connectivity and a moderate to large effect size for the hypo-connectivity alterations, suggesting that these measures may indeed have promise as stratification or treatment biomarkers for ASD.

The above results contribute to a growing consensus on the specific networks characterized by hyper- vs. hypo-connectivity profiles in ASD, but also extend this work to demonstrate a key driver of ASD’s connectivity profile: the larger proportion of frontoparietal voxels within our “DC Increase” mask showing suprathreshold connectivity with voxels outside this mask, and the larger proportion of somatosensory voxels within our “DC Decrease” mask showing subthreshold connectivity with voxels outside this mask. Importantly, our somatosensory findings especially concord with emerging ICA-based analyses of the EU-AIMS LEAP cohort (Oldehinkel et al., in preparation), in which connectivity alterations in ASD are generally most pronounced when considering connectivity between networks found to be distinct in neurotypical subjects. This concordance, despite different preprocessing and analysis methodologies, suggests ample robustness to the results reported here, and motivate further characterization of the etiology, development and clinical symptoms associated with these phenotypes in ASD. At the same time, we note that the most consistent ICA-based findings of between-network differences involve cerebellum, which was only inconsistently acquired in the broader range of cohorts analyzed here. Clearly, the role of cerebellar function in the broader ASD phenotype, and its relation to frontoparietal connectivity in particular, may be an important direction for future work*(31)*.

We do not find consistent age-by-diagnosis interactions on the identified hyper- and hypo-connectivity indices. Nonetheless, descriptively all cohorts showing consistently smaller effect size for the hypo-connectivity index in children suggesting a potential increase of these alterations over age. As this index is not robustly associated with any clinical scales, it may serve a more compensatory role that increases with age. In line with this hypothesis, previous studies reported age-related improvement among older ASD patients in terms of cognitive flexibility*(32)*.

When evaluating the link between imaging and clinical features, we find that the extent of hyper-connectivity abnormalities is correlated with autism symptoms in two out of three cohorts. Hyper-connectivity was associated with social and communication functioning as well as daily living skills. Greater connectivity alterations are thereby associated with greater severity of respective symptoms. These findings further support the idea that these alterations may reflect pathological processes related to skills needed for development of personal independence and social communication abilities. However, the rather low effect size of those associations limits their usability as potential efficacy read-outs. We also failed to replicate this association in the on average six to ten years younger ABIDE II subpopulation. Considering that ABIDE II children diagnosed with ASD do not show any detectable connectivity abnormalities, the lack of significant correlation with clinical symptoms in this cohort is not surprising. Overall, further research is needed to understand the specific reasons for these discrepant findings. Potential confounding effects of medication on functional connectivity have been raised for previous ASD studies *(22)*. Here, we did not find consistent effects of psychotropic medication on the functional connectivity measures with the only significant effect pointing to reduced functional connectivity alterations with psychotropic medication use.

In conclusion, we provide evidence of reproducible hyper- and hypo-connectivity alterations in ASD patients. These alterations are mainly driven by shifts in connectivity from within to outside the respective networks and are linked to core clinical deficits observed in ASD.

## Materials and methods

### Study datasets and replication strategy

To enable an accurate identification and replication of functional connectivity alterations in ASD four datasets were included into the study (Table 1, Supplement 1): EU-AIMS LEAP (www.eu-aims.com), ABIDE I, ABIDE II and InFoR*(9, 33–35)*. All cohorts acquired rs-fMRI data in ASD patients and typically developing volunteers (TD) along with other cohort-specific measures. Only ASD patients with average range IQ (>70) were included for further analyses to reduce variability associated with low functioning ASD*(36)*. In brief, EU-AIMS LEAP cohort is a large well characterized multicenter cohort of ASD patients and TD. ABIDE I and II provide large, retrospectively aggregated multi-center data with varying diagnostic criteria, rs-fMRI sequences and clinic scales. InFoR is a smaller single center cohort with standardized imaging assessments. As the only large cohort with standardized imaging assessments and inclusion/exclusion criteria, the EU-AIMS LEAP dataset was selected for exploratory analyses. The three other cohorts were used for replication. All studies were approved by local ethics committees.

### MRI sequences and rs-fMRI pre-processing

In all cohorts, rs-fMRI data with varying acquisition parameters were acquired. Details on sequences, applied quality criteria and pre-processing are provided in Supplement 1 (Table S1-S4). All pre-processing and analyses were performed using Matlab (R2013b), Statistical Parametric Mapping (SPM12, *(37)*) and IBM SPSS Statistics (Version 23.0). In brief, preprocessing of rs-fMRI data was identical across all cohorts and included normalization into a common space based on structural information, smoothing, and a strict motion correction (s. Supplement 1).

### Identifying functional connectivity alterations

Given the divergent prior findings on functional connectivity alterations in ASD, we followed an unbiased exploratory approach by computing a simple connectivity metric (degree centrality – DC, Supplement 1) to compare ASD patients and TD controls. DC is a count-based measure that assigns to each voxel the sum of all correlation coefficients between the time series of that voxel and all other voxels in the brain exceeding a prespecified threshold (r>0.25). This threshold was recommended in previous studies using DC to eliminate counting voxels that had low temporal correlation attributable to signal noise *(38)*. All further group comparison and correlational analyses were controlled for site, sex, age and IQ effects. As compared to many other connectivity approaches proposed in the literature DC does not require specific assumptions about the location of the signal and can be computed on a voxel-wise basis. Importantly, though dimensionality reduction techniques such as independent component analysis are also often applied in the literature to evaluate functional connectivity, they are less optimal for the key focus of our study which was in replication across different cohorts. Solutions provided by such algorithms are often highly susceptible to differences across study populations such as sample size, demographics and scanner and sequence differences. Also there is no consistent approach for matching the identified components across cohorts. The DC approach was therefore chosen for the replication focus of this study.

The resulting DC maps for EU-AIMS LEAP were entered into a voxel-wise general linear model (GLM) comparing ASD patients and TD using an exact cluster-corrected significance threshold (Supplement 1). Identical GLM designs were then created for ABIDE I, ABIDE II and InFoR data. Weighted mean DC signals (first eigenvariate) were extracted for the three replication cohorts based on EU-AIMS LEAP findings. Effect sizes (Cohen’s *d*) and independent sample *t*-tests were computed comparing DC values between ASD patients and TD (p<.05). We then tested if the whole-brain DC alteration patterns observed in ASD are similar across the cohorts by computing correlations between the un-thresholded *t*-maps (degrees of freedom for p-value computation are based on number of resolution elements in the images provided by SPM12) (Figure 1c, d). As residual motion effects have been repeatedly criticized as potential sources of observed functional connectivity differences between ASD and TD, we have computed mean translational and rotational motion and frame-wise displacement and compared it between ASD and TD using t-tests (Table S12). As the mean frame-wise displacement was indeed significantly higher in ASD as compared to TD, we have also recomputed all analyses by additionally controlling for the effects of the motion parameters in the between group comparison. For this we added mean translational and rotational motion and mean translational and rotational frame-wise displacement for each subject as additional covariates of no interest and recomputed the above analyses.

### Understanding the nature of functional connectivity alterations

Whilst DC provides evidence of altered functional connectivity, it does not allow conclusions on the nature of those alterations. For example, DC changes could arise from alterations in mean strength and/or variance of the underlying connectivity, or shifts in connectivity from within to outside of the respective networks. To better understand the observed functional connectivity alterations, we first computed subject-specific pair-wise Fisher’s *z*-transformed correlations between time courses of all voxels in the brain. Based on these correlations we computed the following four types of connectivity indices within and outside regions showing DC alterations as defined in Figure 2a: 1) mean connectivity of all voxels, 2) variance of connectivity of all voxel-wise correlations, 3) proportion of connected voxels, and 4) shifts in connectivity from inside to outside the respective regions.

Effect sizes and t-tests comparing ASD patients and TD were then computed for each of the above indices. To test if these indices indeed reflect the initial DC changes we computed correlations between both in the EU-AIMS LEAP dataset. Indices showing strongest correlation with initial DC increase and decrease findings and a similar or higher effect size were selected for further evaluations. These two indices providing a more refined view on functional connectivity alterations were extracted for the three replication cohorts and used for further analyses. Both indices are further referred to as hyper- and hypo-connectivity indices.

We further examined in how far the initial choice of the correlational threshold used for DC computation affected the effect sizes observed with both indices. For this we systematically varied in the EU-AIMS LEAP dataset the correlational threshold from −0.25 to 0.7 computing the effect sizes for differentiation between ASD and TD.

### Understanding the link between imaging findings and clinical and demographic factors

Having identified the hyper- and hypo-connectivity indices as replicable ASD biomarkers we next investigated their relationship to clinical symptoms, medication status, age and sex. To test for the effects of age, we split the cohorts into three age categories: children (below 12), adolescents (age between 12 and 18) and adults (age above 18). Hyper- and hypo-connectivity indices between ASD patient and respective TD populations were compared by computing effect sizes and t-tests. Additionally, we formally tested for age-by-diagnosis interactions (three age categories) using ANOVAs controlling for sex, site and IQ. Similarly, to test for the effects of sex we computed ANOVAs testing for sex-by-diagnosis interactions controlling for age, site and IQ. We further evaluated for each cohort if psychotropic medication interferes with the observed functional connectivity alterations. Because detailed medication information on specific drugs, dose or duration was not available we restricted the analyses to t-tests comparing DC and hyper- and hypo-connectivity indices between patients on and off psychotropic medication.

To test for potential relationships between hyper- and hypo-connectivity indices and clinical scales, we followed a similar exploration and replication strategy as for the group comparisons. Because of few overlaps regarding clinical scales and a much smaller number of patients than in the exploration sample InFoR was not used for these analyses. Separate GLMs (as implemented in SPSS) were computed using the hyper- and hypo-connectivity indices to predict clinical severity in the EU-AIMS LEAP ASD patients (Table S10). Significant (*p*<0.05) or marginally significant (*p*<0.1) relationships identified in the EU-AIMS cohort were selected for replication in ABIDE I and II using identical GLMs.

## Acknowledgements

EU-AIMS LEAP is supported by the Innovative Medicines Initiative Joint Undertaking under grant agreement number 115300 (EU-AIMS), resources of which are composed of financial contribution from the European Union’s Seventh Framework Programme (FP7/2007 - 2013) and the European Federation of Pharmaceutical Industries and Associations (EFPIA) companies’ in kind contribution.

ABIDE I is supported by NIMH (K23MH087770), NIMH (R03MH096321), Leon Levy Foundation, Joseph P. Healy and the Stavros Niarchos Foundation.

ABIDE II is supported by NIMH (5R21MH107045), NIMH (5R21MH107045), Nathan S. Kline Institute of Psychiatric Research, Joseph P. Healey, Phyllis Green and Randolph Cowen.

InFoR is part of clinical trial C07-33 sponsored by Inserm. It was granted approval by local Ethics Committee or “Comité de Protection des Personnes” on 2008 November 14th, authorized by the French authorities (ANSM B80738-70 on 2008 August 11th), and registered in a public trials registry (NCT02628808). All study participants gave their informed written consent to participation, in line with French ethical guidelines. The work has been co-ordinated by Fondation FondaMental and achieved thanks to the following organisms and establishments: AP-HP, CHU Bordeaux, Hôpital Charles Perrens, Robert Debré et Henri Mondor’s CIC. It was financially supported in part by the Institut Roche, in part by the Investissements d’Avenir program managed by the ANR under reference ANR-11-IDEX-0004-02.

## Authors’ contributions

SH and JD designed the overall study, performed the analyses and wrote the manuscript. AB, GDH, FB, TC, JT, CFB, EL, DM, MO, CB, AR, JKB, WS, PG, JFH, JS and CHC contributed to design, collection and interpretation of the EU-AIMS data. CC, MD, ML, IS, AG, RD, AB, CL, JH, MLL, SG, MC, AA, MB, CB, XL and AnB contributed to design, collection and interpretation of the InFoR data. All authors reviewed and commented on the manuscript.

## List of Supplementary Materials

Supplement 1

